# Catalytic antibodies in arrhythmogenic cardiomyopathy patients cleave desmoglein 2 and N-cadherin and impair cardiomyocyte cohesion

**DOI:** 10.1101/2023.02.08.527624

**Authors:** Sunil Yeruva, Konstanze Stangner, Anna Jungwirth, Matthias Hiermaier, Maria Shoykhet, Daniela Kugelmann, Michael Hertl, Shohei Egami, Norito Ishii, Hiroshi Koga, Takashi Hashimoto, Michael Weis, Britt Maria Beckmann, Ruth Biller, Dominik Schüttler, Stefan Kääb, Jens Waschke

**Affiliations:** Chair of Vegetative Anatomy, Faculty of Medicine, Ludwig-Maximilians-University (LMU) Munich, Munich, Germany; Department of Dermatology and Allergology, Philipps-University Marburg, Marburg, 35037, Germany; Department of Dermatology, Keio University School of Medicine, Tokyo, Japan; Department of Dermatology, Kurume University School of Medicine, Kurume, Japan; Department of Dermatology, Osaka City Metropolitan University, Graduate School of Medicine, Osaka, Japan; Krankenhaus Neuwittelsbach, Fachklinik für Innere Medizin, Munich, Germany; Department of Medicine I, University Hospital Munich, Campus Großhadern, Ludwig-Maximilians University Munich (LMU), Munich, Germany; Institute of Legal Medicine, University Hospital Frankfurt, Goethe University, Frankfurt, Germany; ARVC-Selbsthilfe e.V., Patient Association, Munich, Germany; DZHK (German Centre for Cardiovascular Research), Partner Site Munich, Munich Heart Alliance (MHA), Munich, Germany; Walter Brendel Centre of Experimental Medicine, Ludwig-Maximilians University Munich (LMU), Munich, Germany; Interfaculty Center for Endocrine and Cardiovascular Disease Network Modelling and Clinical Transfer (ICON), LMU Munich, Munich, Germany

**Keywords:** Arrhythmogenic cardiomyopathy, Intercalated disc, Intercellular adhesion, Autoantibodies, Desmoglein 2, Cadherin, Pemphigus, Desmosome

## Abstract

**Aims:** Arrhythmogenic cardiomyopathy (AC) is a severe heart disease predisposing to ventricular arrhythmias and sudden cardiac death caused by mutations affecting intercalated disc (ICD) proteins and aggravated by physical exercise. Recently, autoantibodies targeting ICD proteins, including the desmosomal cadherin desmoglein 2 (DSG2), were reported in AC patients and were considered relevant for disease development and progression, particularly in patients without underlying pathogenic mutations. However, it is unclear at present whether these autoantibodies are pathogenic and by which mechanisms show specificity for DSG2 and thus can be used as a diagnostic tool.

**Methods and Results:** IgG fractions were purified from 15 AC patients and 4 healthy controls. Immunostainings dissociation assays, atomic force microscopy (AFM), western blot analysis and Triton-X-100 assays were performed utilizing human heart left ventricle tissue, HL-1 cells, and murine cardiac slices. Immunostainings revealed that autoantibodies against ICD proteins are prevalent in AC and most autoantibody fractions have catalytic properties and cleave the ICD adhesion molecules DSG2 and N-cadherin, thereby reducing cadherin interactions as revealed by AFM. Furthermore, most of the AC-IgG fractions causing loss of cardiomyocyte cohesion activated p38MAPK, which is known to contribute to a loss of desmosomal adhesion in different cell types, including cardiomyocytes. In addition, p38MAPK inhibition rescued the loss of cardiomyocyte cohesion induced by AC-IgGs.

**Conclusion:** Our study demonstrates that catalytic autoantibodies play a pathogenic role by cleaving ICD cadherins and thereby reducing cardiomyocyte cohesion by a mechanism involving p38MAPK activation. Finally, we conclude that DSG2 cleavage by autoantibodies could be used as a diagnostic tool for AC.

## Introduction

Arrhythmogenic cardiomyopathy (AC) is a rare genetic heart disease potentially causing sudden cardiac death (SCD), especially among athletic young adults ^1, 2^. The prevalence of this disease had been estimated between 1:1000 to 1:5000 ^3, 4^. Of note, SCD can occur even during the first concealed phase without any observable structural changes of the ventricles and SCD is the first disease manifestation in about 5-10 % of cases ^3, 5^.

AC is a familial disease, and 60 % of patients carry mutations in genes coding for desmosomal proteins of the ICD ^6, 7^. ICDs are specialized intercellular contact areas, wherein desmosomes and adherens junctions (AJ) (providing mechanical adhesion), together with gap junctions (GJ) and sodium channels (important for electric coupling), form a complex structure referred to as the connexome ^8^. Within the connexome, excitation propagation is mediated by GJ via electrotonic and ephaptic coupling with Nav1.5 sodium channels ^9^. In AC patients, mutations in genes coding for desmosomal proteins, include both the plaque proteins, such as plakophilin 2 (*PKP2*), desmoplakin (*DSP*) and plakoglobin (*JUP*), and the cadherin-type adhesion molecules, such as desmoglein 2 (*DSG2*) and desmocollin 2 (*DSC2*). Gene mutations for N-cadherin (*N-CAD*), desmin (*DES*), transmembrane protein 43 (*TMEM43*), sodium voltage-gated channel subunit 5 (*SCN5A or Nav1.5*), were rarely found ^10–12^. Interestingly, the clinical phenotype differs with respect to the underlying mutation and patients with *DSP* mutations often exhibit a left dominant or biventricular involvement with left ventricular failure. In contrast, patients with other gene mutations such as *PKP2* mostly suffer from the predominant right-ventricular disease ^13^. Therefore, DSP cardiomyopathy (DPC) is clinically distinguished from typical arrhythmogenic right ventricular cardiomyopathy (ARVC).

However, no genetic cause can be determined for a substantial number of patients, and the course of the disease can differ significantly in families with the same underlying mutation, thus the genetic predisposition alone cannot explain the pathogenesis of AC. In this context, it is highly relevant that autoantibodies targeting DSG2 have been described in AC patients, irrespective of the AC causing gene mutations ^14^. Anti-DSG2 antibodies therefore might be useful for future diagnosis of AC since they were reported to be specific for AC ^14^. In addition, a recent study performed in a large cohort of AC patients found that the majority of familial and sporadic AC patients had antibodies against heart and ICD proteins ^15^.

Nevertheless, these studies did not evaluate whether these autoantibodies can directly or indirectly affect the mechanical strengths of the ICDs and therefore are pathogenic. Thus, AC pathogenesis may have aspects similar to pemphigus vulgaris (PV) – an autoimmune blistering skin disease caused by autoantibodies against DSG 3 ^16–18^. This is of particular interest since both the structure of the ICD and the mechanisms regulating intercellular adhesion in cardiomyocytes share similarities to desmosomes in keratinocytes ^19–21^.

In this study, we demonstrate for the first time that IgGs in AC patients exhibit catalytic activity against DSG2 and N-CAD effectively reducing interaction of these adhesion molecules. Furthermore, most AC-IgGs that caused loss of cardiomyocyte cohesion induce activation of p38MAPK, whereas inhibition of the p38MAPK restored the loss of cardiomyocyte cohesion caused by the former.

## Results

### AC Patient details

This study included blood samples from 15 AC patients (AC1-15) from 8 different families, as well as 4 healthy unrelated controls. Among the AC patients, 11 were males with an average age of 41±4.5 years (mean ± SEM), and 4 were females with an average age of 46±9.5 years (mean ± SEM). We divided the AC patients into two groups, DPC carrying *DSP* gene mutations (9 patients from 2 different families) and arrhythmogenic right ventricular cardiomyopathy (ARVC) carrying *PKP2* gene mutations (6 patients from 6 different families) as per the recent report ^22^. Pooled isolated IgGs from 4 healthy donors (2 males and 2 females) served as a control group (referred as IgG in the figure panels). 6 of the 9 DPC patients and 6 of the 6 ARVC patients met the task force criteria of 2010 ^6, 23^. Detailed characteristics of the patients’ clinical features are given in Table 1.

**Table 1:** Arrhythmogenic cardiomyopathy (AC) patient details

| **Gene Mutation** | **Patient ID (Family)** | **Task force criteria** | **Age** | **Sex** | **Clinics** |
| --- | --- | --- | --- | --- | --- |
| *DP*<br>(*p. Gln952\**) | AC1<br>(1*) | Yes | 59 | male | PPK, supraventricular arrhythmia, RV enlargement and + WMA, LVEF 50 % |
| *DP*<br>(*p. Gln952\**) | AC2<br>(1*) | Yes | 33 | male | syncope, RV+LV enlarged + dysfunction, LGE RV and LV, LVEF 50 %, RVEF 49 % |
| *DP*<br>(*p.Arg 1934\**), | AC4<br>(2) | Yes | 34 | male | PPK, curly hair, premature ventricular contractions, LVEF 54 %, RVEF 61 %, LGE LV |
| *DP*<br>(*p. Gln952\**) | AC7<br>(1*) | Yes | 27 | male | ventricular tachycardia/ventricular fibrillation, LVEF 64 %, normal RV function |
| *DP*<br>(*p. Gln952\**) | AC8<br>(1*) | No | 34 | male | asymptomatic, normal RV, LV function |
| *DP*<br>(*p. Gln952\**) | AC9<br>(1*) | Yes | 20 | male | supraventricular arrhythmia, RV enlargement+ WMA, LVEF 62 %, reduced RVEF |
| *DP*<br>(*p. Gln952\**) | AC10<br>(1*) | Yes | 31 | male | palpitations, LVEF 51 %, LGE LV, RVEF 42 % |
| *DP*<br>(*p. Gln952\**) | AC12<br>(1*) | No | 64 | fem. | polymorphic premature ventricular contractions, LVEF 45 %, normal RV function |
| *DP*<br>(*p.Arg 1934\**), | AC13<br>(2) | No | 30 | fem. | PPK, curly hair, arrhythmias, premature ventricular contractions/ventricular tachycardia, LV dysfunction, LVEF 30-35%, RV LGE, RVEF normal |
| *PKP2*(*p. Arg79\**) | AC3<br>(3) | Yes | 61 | fem. | fibrofatty dysplasia in biopsy, supraventricular arrhythmias, ventricular tachycardia |
| *PKP2* | AC5<br>(4) | Yes | 41 | male | ventricular tachycardia, RV dilation, RV dysfunction, LVEF 60% |
| *PKP2* | AC6<br>(5) | Yes | 57 | male | ventricular arrhythmia, RV enlargement, RV dysfunction, RVEF 45%, LVEF 55% |
| *PKP2* | AC11<br>(6) | Yes | 29 | fem. | ventricular arrhythmia (ventricular tachycardia, ventricular fibrillation), atrial fibrillation, RV enlargement + WMA, LVEF 60% |
| *PKP2 (1688 +1 G>A)* | AC14<br>(7) | Yes | 47 | male | ventricular tachycardia, RV enlargement and dysfunction+ WMA, RVEF 19%, LVEF 59%, heart transplantation 2020 |
| *PKP2 (c.369G>A)* | AC15<br>(8) | Yes | 68 | male | , ventricular tachycardia, RV enlargement, severe RV dysfunction, RVEF 18 %, right ventricular assist device, LVEF 54 % |

DSP: desmoplakin, LGE: late gadolinium enhancement, LV: left ventricle, LVEF: left ventricular ejection fraction, PKP2: plakophilin 2, PPK: palmoplantar keratoderma, RV: right ventricle, RVEF: right ventricular ejection fraction, WMA: wall motion abnormality.

**Table 2:** Brief summary of the important findings of this study.

| AC sample | Immunostaining ICD | Cleavage/AFM DSG2 | Dissociation assay |
| --- | --- | --- | --- |
| **DPC** |  |  |  |
| Control IgG | - | -/- | - |
| AC1 | + | +/+ | + |
| AC2 | + | +/+ | + |
| AC4 | + | -/n.a. | - |
| AC7 | + | +/+ | + |
| AC8 | + | +/n.a. | - |
| AC9 | + | +/n.a. | + |
| AC10 | + | +/n.a. | - |
| AC12 | + | +/n.a. | - |
| AC13 | + | +/+ | - |
| **ARVC** |  |  |  |
| AC3 | + | -/n.a. | - |
| AC5 | + | -/+ | + |
| AC6 | + | +/+ | - |
| AC11 | + | +/+ | - |
| AC14 | + | +/n.a. | - |
| AC15 | + | +/n.a. | + |

### AC patients carry antibodies against ICD proteins

Autoimmunity is frequently involved in different cardiovascular diseases such as heart failure, myocarditis, dilated and ischemic cardiomyopathies ^24, 25^. Two recent studies using a large cohort of AC patients demonstrated the presence of autoantibodies either against anti-heart and anti-ICD proteins ^15^ or specifically against the desmosomal cadherin protein, DSG2 ^14^. We further evaluated the presence of autoantibodies targeting ICD proteins in our patient samples. Therefore, we examined AC patient IgGs by immunostaining utilizing human heart ventricular tissue to assess the presence of anti-ICD antibodies. N-CAD was used as a marker for ICDs. As a positive control for DSG2 a commercial human anti-DSG2 antibody was used. All AC-IgGs (AC1-AC15) showed positive staining of ICDs with varying intensities (Fig. 1). In contrast, control IgGs from the four healthy individuals showed no ICD staining (Fig. 1) and goat anti-human IgG secondary antibody did not show cross-reactivity with ICDs (Supplementary fig. 1).

**Fig. 1.**
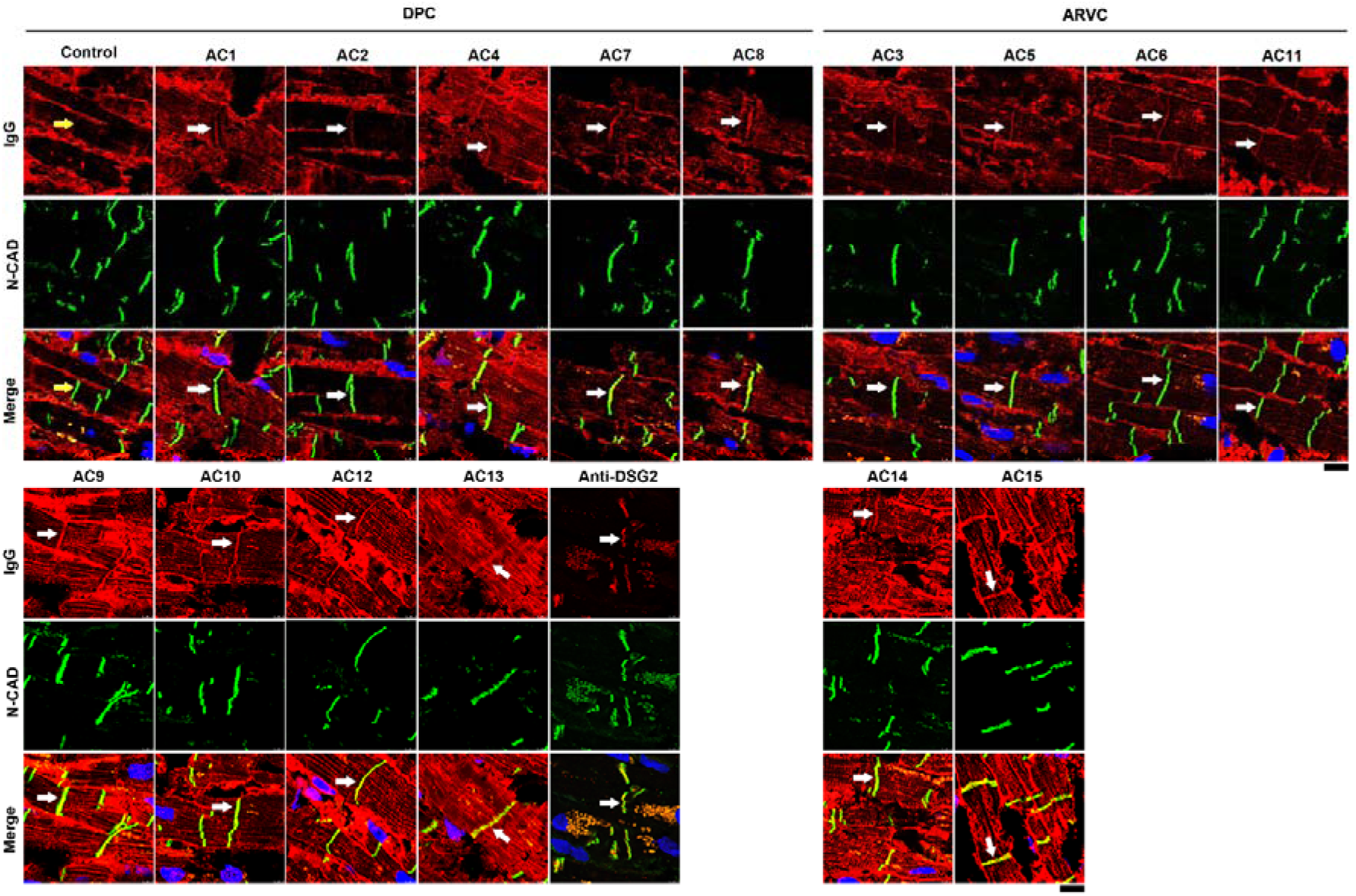
Detection of antibodies against ICD proteins in human left ventricular tissue by immunofluorescence analysis. Human left ventricular tissue was incubated with respective AC-IgGs and Control-IgGs as indicated in the figure. N-CAD was used as a marker for ICDs, and presence of anti-ICD proteins was defined when there was an overlap of IgG staining with N-CAD (white arrows). The yellow arrow in control-IgG shows no positive staining of ICDs. Anti-DSG2 antibody was used to show the presence of DSG2 in the cardiac tissue used. AC patients were grouped into DPC (harboring desmoplakin gene mutations) and ARVC (harboring plakophilin2 gene mutations). Images represent immunofluorescence analysis of 3 repeats. Scale bar: 10 µm.

### AC autoantibodies disturbed cardiomyocyte cohesion in HL-1 cells and murine cardiac slice culture

Since we detected anti-ICD antibodies among AC patient IgGs, we hypothesized that autoantibodies in AC patients might cause alterations in cardiomyocyte cohesion. We utilized HL-1 cells, a murine atrial cardiomyocyte cell line, and performed dissociation assays (Fig. 2A). As a positive control, we used a commercial anti-DSG2 antibody targeting the extracellular domain of human DSG2 to show that antibodies targeting human DSG2 can impair cardiomyocyte cohesion. In a well-established dissociation assay ^26, 27^ incubation with 6 out of 15 AC-IgG fractions for 24 hours resulted in a significant reduction in HL-1 cell cohesion (Fig. 2A-C; Supplementary fig. 2A and B). AC patients whose IgGs caused a loss of cohesion were males and met the task force criteria for AC. However, not all IgG fractions from patients with task force criteria induced loss of cardiomyocyte cohesion. We further investigated whether results from HL-1 cell line can be replicated in cardiac tissue. For this purpose, we performed dissociation assays utilizing murine cardiac slices and treated them with one patient IgG from each group (AC1 from DPC group and AC15 from ARVC group). After 6-hour incubation of the respective AC-IgGs, we detected a significant loss of cardiomyocyte cohesion in the murine cardiac slices compared to the healthy control IgG (Fig. 2D-F).

**Fig. 2.**
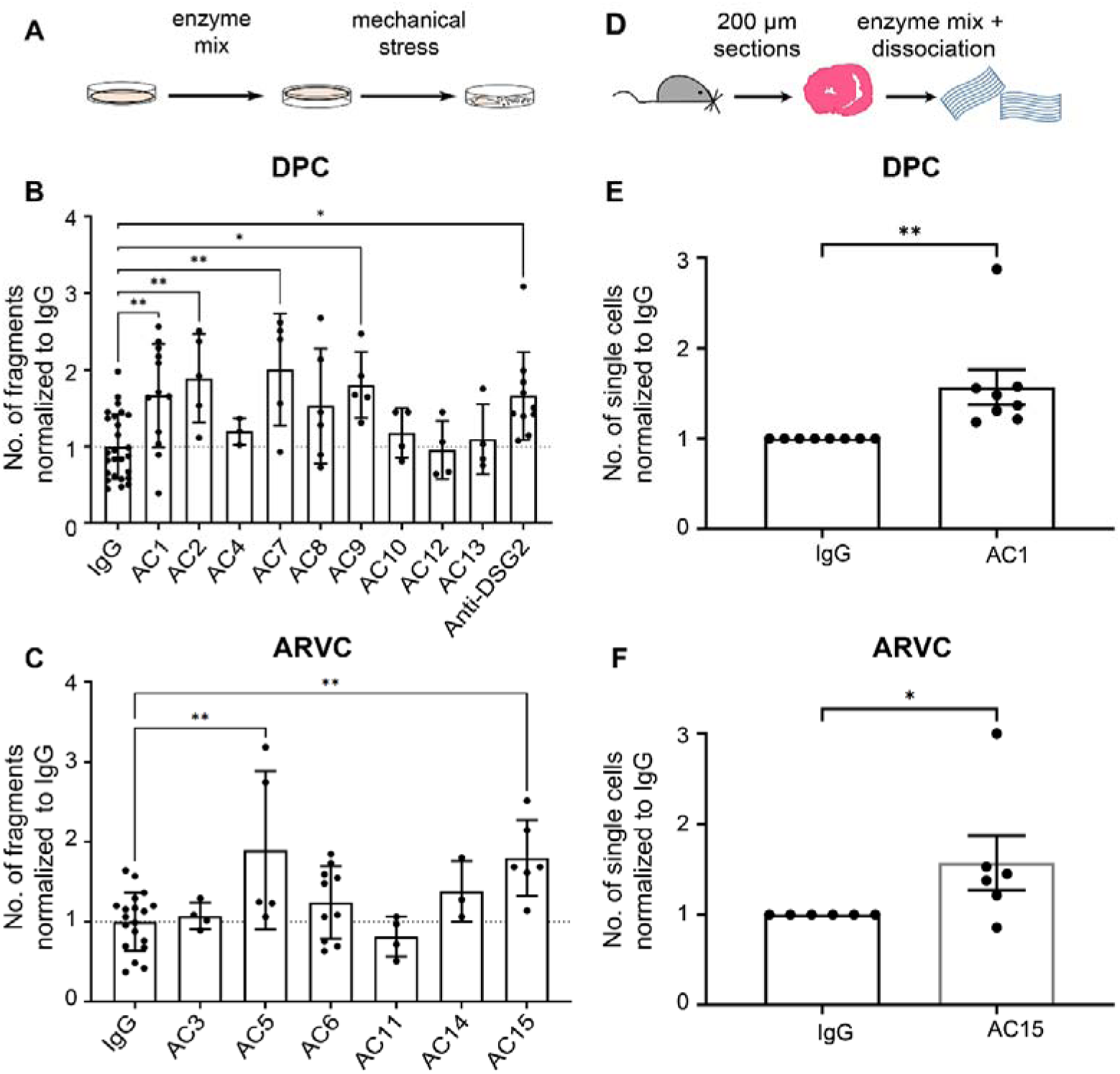
Dissociation assays in HL-1 cells and murine cardiac slices **A**. Dissociation assays were performed in HL-1 cells (a cartoon illustrates the method) treated with **B**. DPC patient IgGs and **C**. ARVC patient IgGs for 24 hours. The increase in the number of fragments correlates to a decrease in cell cohesion and vice versa. In each experiment, duplicate or triplicate wells were incubated with respective IgGs, and at least a minimum of 3 biological repeats were performed. Bar graph represents fold change compared to healthy control-IgG (IgG). Data are presented as mean ± SEM. One-way ANOVA Holm-Šídák’s multiple comparisons test was performed. **D.** Dissociation assay in murine cardiac slice cultures (a. cartoon illustrates the method) obtained from wild-type mice treated with **E.** AC1 (from DPC group as in **B**) and **F.** AC15 (from ARVC group as in **C**). N=6-8 mice, Bar graph represents fold change compared to healthy control IgGs (IgG). Data are presented as mean ± SEM. Paired *t-test*. * Indicates p<0.05 ** indicates p<0.005.

### AC patients IgGs exhibit catalytical properties to cleave DSG2 and N-CAD

As autoantibodies against DSG2 were recently detected in AC patients ^14^, we further investigated whether the observed loss of cardiomyocyte cohesion induced by some AC-IgGs was caused by autoantibodies targeting DSG2. We thus performed a DSG2-specific ELISA analysis utilizing human DSG2 protein tagged with His. Although the positive control, which detects the extracellular region of DSG2, showed a strong signal, none of the tested AC patient IgG fractions showed a signal beyond the cut-off OD value (Supplementary fig. 3A). Since in a previous study by Chatterjee et al., DSG2 protein tagged with Fc (DSG2-Fc) was used to detect antibodies specific to DSG2 using ELISA analysis ^14^, we also performed ELISA adopting the same approach. However, the secondary goat anti-human antibody alone (used as negative control) showed a similar OD as all AC-IgG fractions, and the controls. In addition, these results were comparable to two different monoclonal DSG2 antibodies (Supplementary fig. 3B). Therefore, we adopted the Western blot technique as performed by Chatterjee et al. ^14^. We used DSG2-Fc protein and N-CAD-Fc as another ICD protein and performed Western blots. Since we used DSG2 and N-CAD tagged with Fc fragments, the goat-anti-human IgG-horseradish peroxidase (IgG-HRP) antibody was diluted to a level where no signal was detected (data not shown). However, incubation of blots with respective IgGs revealed both DSG2 and N-CAD protein bands (Supplementary fig. 3C and D). Similarly, incubation with control-IgGs from healthy donors detected the same bands as AC-IgGs, whereas secondary antibody alone did not. Thus, we conclude that the presence of the Fc tag to the proteins might mask the detection sensitivity of the Western blotting and ELISA and therefore are not suitable for detection of autoreactive antibodies.

Since we observed a loss of cardiomyocyte cohesion but did not detect autoantibodies firmly binding to DSG2 in the AC patient IgG fractions, we wondered whether autoantibodies affecting cell-cell adhesion target adhesion molecules by modifying the epitope structure which would interfere with detection by ELISA or Western blot analysis. The presence of catalytic antibodies, which act like proteases, are well known in autoimmune and viral disorders ^28–33^. To evaluate this hypothesis, we developed a cleavage assay in which the IgG fractions of AC patients’ and control were incubated with or without protease inhibitor and DSG2-Fc protein. The resulting fragments were then analyzed by Western blot (Fig. 3A). We detected a 100 kDa DSG2 fragment after incubation with 11 out of 15 patient IgG fractions. The patient IgG fractions resulting in a 100 kDa DSG2 fragment belonged to 5 different families. In contrast, after incubation with control IgG or water, we did not detect the 100 kDa DSG2 fragment. No cleavage was detectable after incubation with AC3-IgG, AC4-IgG, AC5-IgG and AC11-IgG, all of which belonged to separate families. Among those, only AC5-IgG reduced cardiomyocyte cohesion. However, in the presence of a protease inhibitor, no cleavage product was detected in any of the samples.

**Fig. 3.**
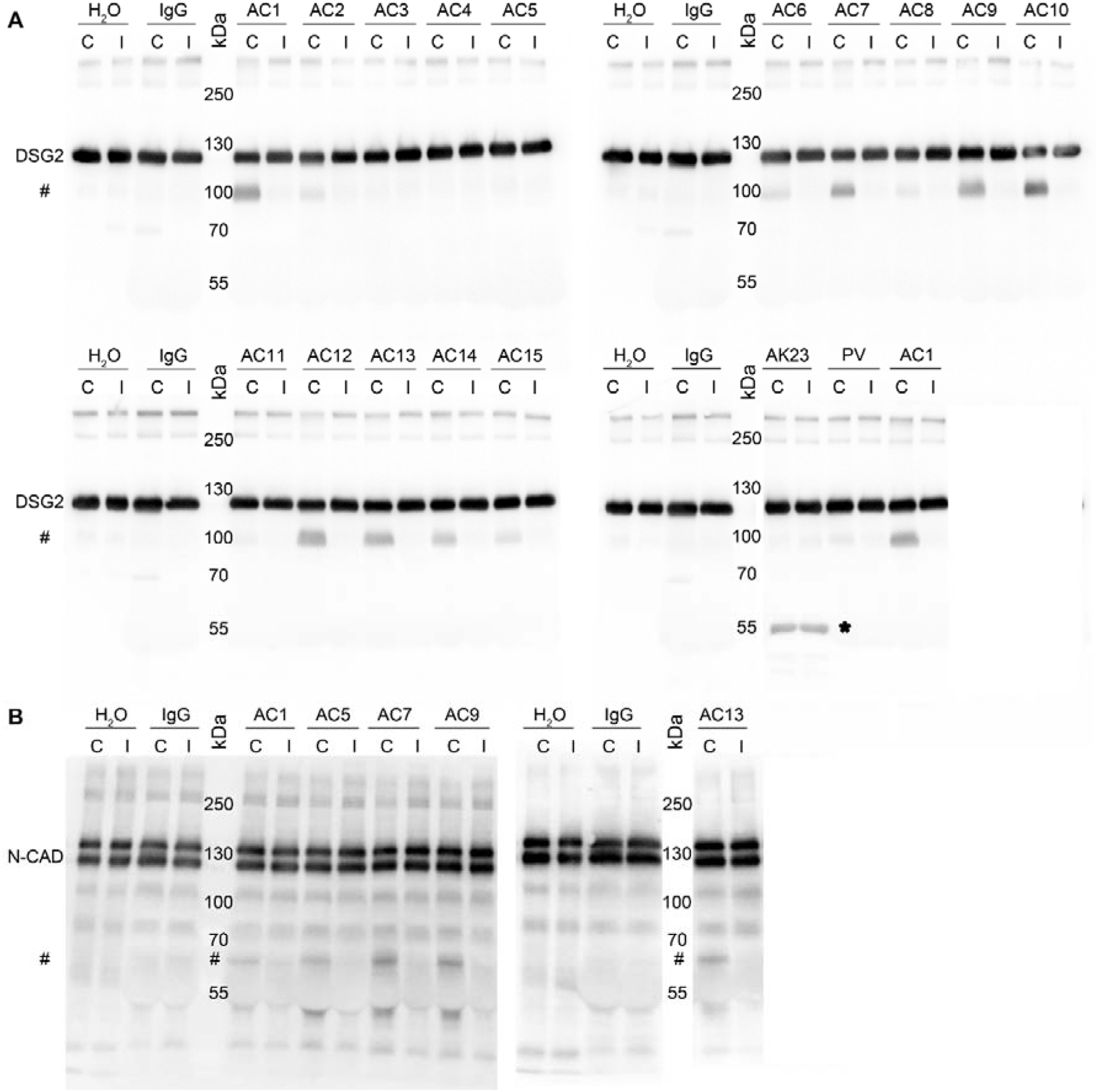
*In vitro* DSG2 and N-CAD Cleavage assays **A.** Human DSG2-Fc protein (200ng/lane) or **B.** N-CAD-Fc protein (200 ng/lane) was incubated with respective IgGs from AC patients (AC1-AC15), water (H_2_O), healthy control IgG (IgG), PV-IgG and AK23 (murine pemphigus monoclonal anti-DSG3 antibody, as a negative control) for 4 hours, without (C) and with protease inhibitor (I, cOmplete™) and Western blot analysis was performed. A representative image of 3 experimental repeats is shown. # Indicates the cleaved fragments. * Indicates mouse IgG heavy chain.

To further evaluate if the observed DSG2 cleavage is specific for autoantibodies from AC patients, we tested whether autoantibodies causing the autoimmune blistering skin disease pemphigus vulgaris (PV), which are directed primarily against DSG3, also induce DSG2 cleavage. We used AK23, a DSG3-specific autoantibody derived from a pemphigus mouse model ^34^ as well as IgGs from a patient suffering from PV. PV-IgG caused a very mild DSG2 cleavage whereas AK23 did not. Moreover, we used a human Fc-construct for DSG3, which is expressed in cells from stratified epithelia such as the epidermis, and performed the same assay by incubating control IgG, AK23, PV-IgG or AC1-IgG, respectively (Supplementary fig. 4). We did not detect DSG3-Fc cleavage after incubation with either pemphigus autoantibodies or AC1-IgG. These results confirm that autoantibodies from AC patients are specific for cleaving DSG2 and that catalytic antibodies against DSG3 are absent in pemphigus.

It was proposed that autoantibodies in AC are specific for DSG2 ^14^. Besides desmosomes, AJs provide mechanical strength to the ICD and thus autoantibodies against N-CAD also could contribute to a loss of cardiomyocyte cohesion. Therefore, we used our cleavage assay to test if AC-IgG fractions were effective to cleave N-CAD. We tested AC-IgGs which reduced cardiomyocyte cohesion in dissociation assays and AC13 IgG as it had no significant effect on cardiomyocyte cohesion. All tested AC-IgGs cleaved N-CAD, which resulted in an approximately 65 kDa fragment. Interestingly, AC5-IgG which impaired cardiomyocyte cohesion in dissociation assay but did not cleave DSG2, was effective to cleave N-CAD. Again, incubation with either water or control IgGs led to no cleavage products. These data indicate that some AC patient IgGs include catalytic antibodies against both DSG2 and N-CAD.

### AC patient IgGs reduced homophilic DSG2 and N-CAD interactions

As we detected the presence of catalytic antibodies in AC patient IgGs, we further asked whether the catalytic properties of these antibodies have any role in disrupting DSG2 or N-CAD binding on a molecular level. Therefore, we performed cell-free atomic force microscopy (AFM) to study the role of AC IgGs in disrupting homophilic DSG2 and N-CAD binding as described before ^26^. In this experiment, a cantilever with DSG2-Fc or N-CAD-Fc-coated tip was approached and used to probe a mica sheet coated with the same protein construct to investigate interaction properties (Fig. 4A). Upon treatment with the respective IgGs, the binding frequency of homophilic DSG2 (Fig. 4B and C) and N-CAD (Fig. 4D) interactions significantly decreased for IgGs from AC1, AC7 and AC13, compared with the healthy control-IgG, whereas AC2 reduced only DSG2 binding frequency. The binding forces were not affected significantly (data not shown). These data indicate that homophilic DSG2 or N-CAD interaction reduced by some AC-IgGs (AC1, AC2, AC5, AC7) correlated with loss of cardiomyocyte cohesion, whereas other AC-IgGs (AC6, AC11 and AC13) caused loss of DSG2 interaction on the molecular level but were not effective to reduce cardiomyocyte cohesion.

**Fig. 4.**
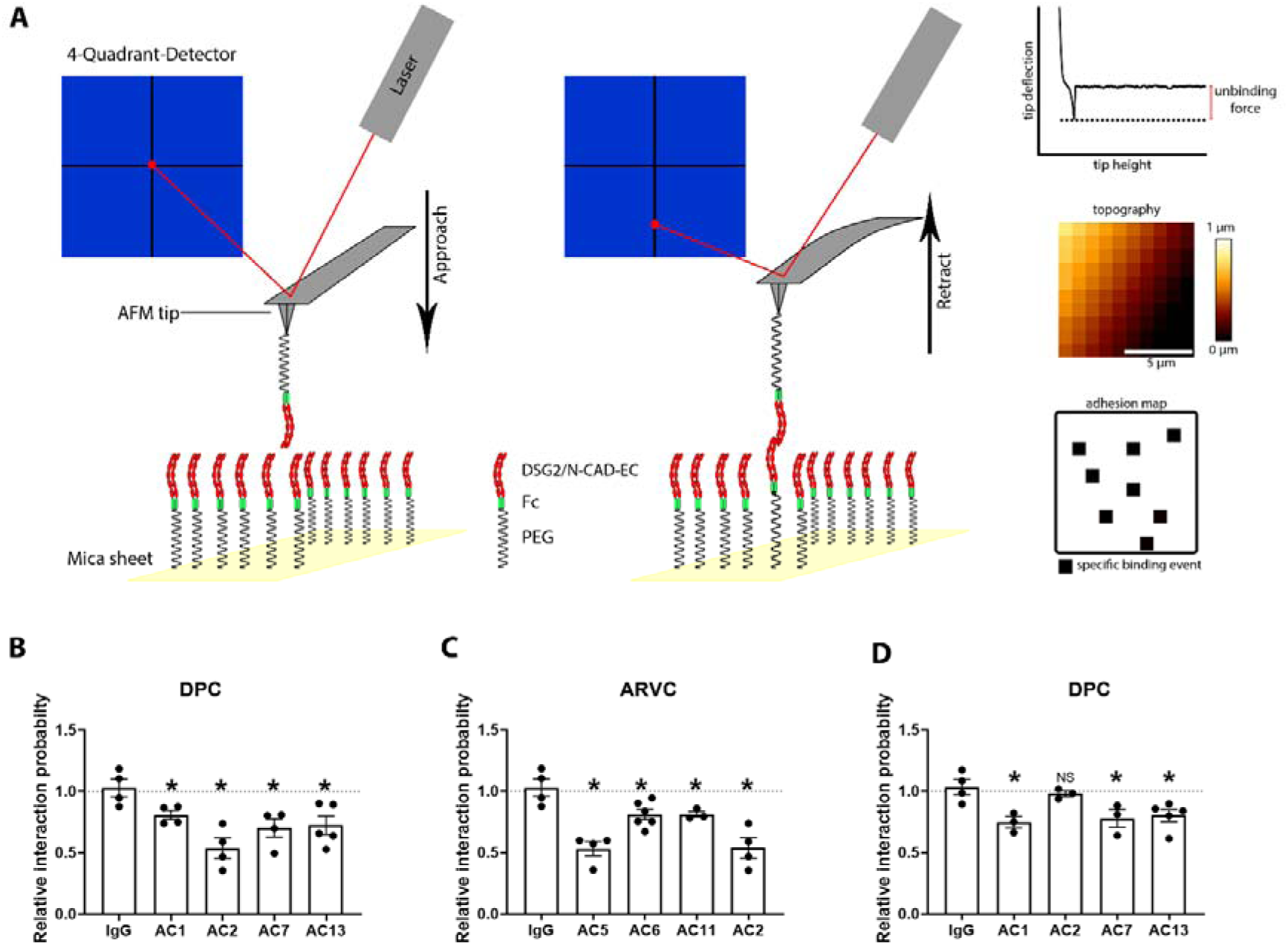
Homophilic DSG2 and N-CAD interaction probabilities measured by AFM **A**. Schematic presentation of an AFM interaction experiment in cell-free conditions. Recombinant DSG2/N-CAD extracellular domain containing proteins tagged with Fc fragments were covalently linked via a PEG linker to the AFM tip and the mica sheet. A laser is directed to the cantilever tip and reflected onto a photodetector. Each interaction event leads to a deflection of the cantilever, detected by the laser reflected on the photodetector and a force-distance curve of a specific binding event was produced, together with a topography image and adhesion events. Quantification of DSG2 binding frequency expressed in relative interaction probability. **B**. in desmoplakin mutated AC patients (DPC), **C**. plakophilin 2 mutated patients (ARVC). **D**. Quantification of N-CAD binding frequency expressed in relative interaction probability in DPC. Each data point represents 1000 curves analyzed across two areas (10 µm x 10 µm). Data are presented as mean ± SEM.* p ≤ 0.05 compared to IgG, and NS is not significant compared to IgG. One-way ANOVA with Holm-Šídák’s multiple comparisons test was performed. N=3-5.

### Pathogenic AC-IgG caused p38MAPK activation

Since about one-third (6 out of 15) AC-IgG fractions caused loss of cardiomyocyte cohesion, we tested whether p38MAPK activation was induced similar to anti-desmoglein autoantibodies (PV-IgGs) in pemphigus ^17, 35–37^. Western blot analysis revealed that AC7-IgG, AC8-IgG, and AC9-IgG caused p38 MAPK activation (Fig. 5A and B, supplementary fig. 5A). Similarly, we performed Triton-X-100 protein extraction following incubation with AC2-IgG and AC5-IgG, which revealed significant p38MAPK activation in the non-cytoskeleton-bound fraction for AC5-IgG (Fig. 5C and D, Supplementary fig. 5 B). AC2-IgG also enhanced p38MAPK activity in Western blot analysis but due to expression variability, quantification of results showed a trend which was not significant. Taken together, 5 AC-IgG fractions, all of which caused DSG2 cleavage and/or loss of DSG2 binding, induced p38MAPK activation. Moreover, except AC8-IgG all AC-IgGs caused a loss of cardiomyocyte cohesion. However, no changes were observed for the distribution of either desmosomal proteins and N-CAD or Cx43 in Western blot analysis and triton assays (Fig. 5A and C and Supplementary fig. 5A and B). To evaluate whether activated p38MAPK contributes to the loss of cardiomyocyte cohesion caused by AC patient IgGs, we performed dissociation assays in the presence of the p38MAPK inhibitor SB202190 (SB20) utilizing HL-1 cells. Dissociation assays revealed that an inhibition of p38MAPK rescued the loss of cardiomyocyte cohesion caused by AC-IgGs (Fig. 5E and Supplementary fig. 5C).

**Fig 5:**
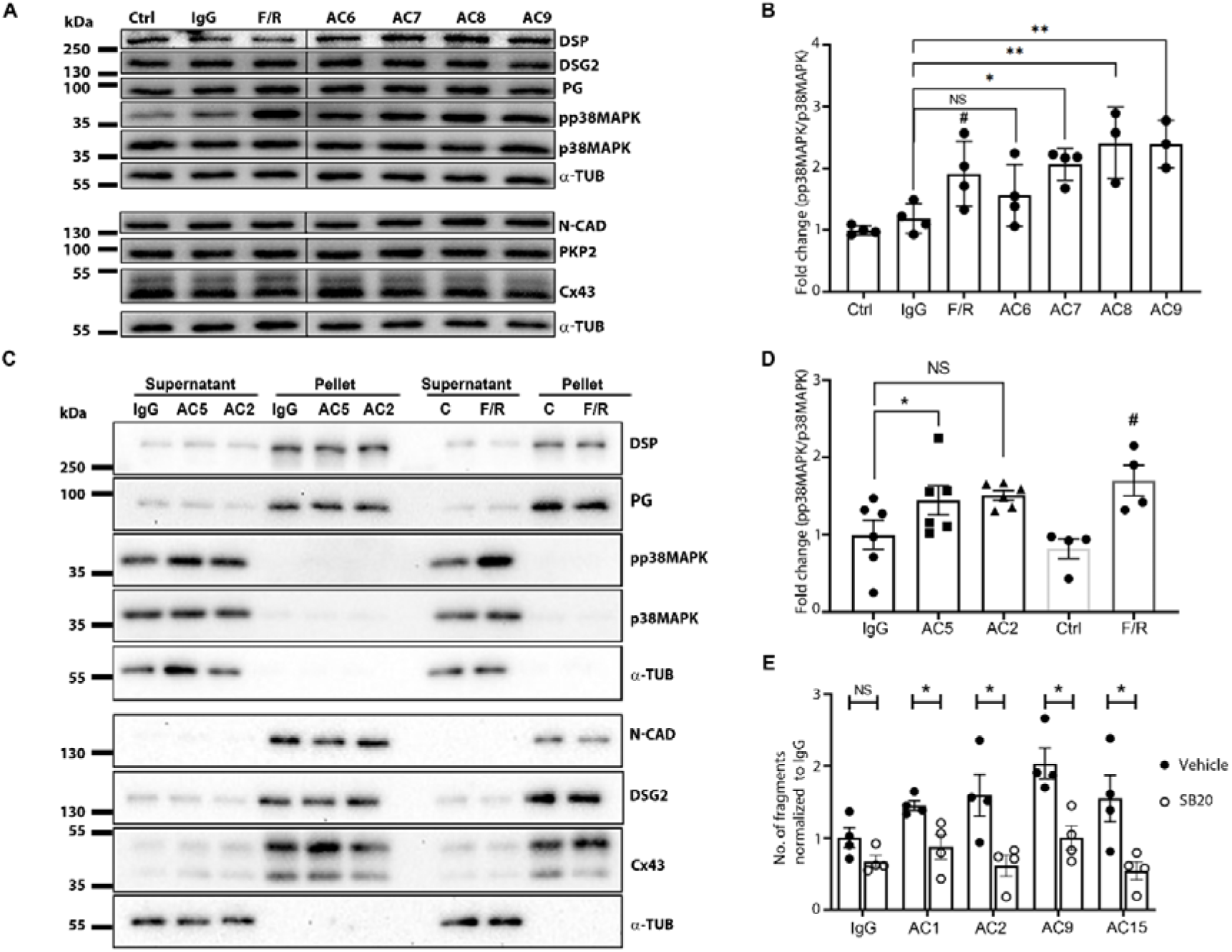
Protein expression analysis and dissociation assays in HL-1 cells treated with AC IgGs **A.** Western blot of lysates prepared from HL-1 cells treated with respective IgGs for 24 hours. **B.** Western blot quantification of pp38MAPK for the blots shown in A. Ratio between pp38MAPK and total p38MAPK was normalized to healthy control IgG (IgG) and means represented ± SEM. N=3–4. **C**. Western blots of Triton-X-100 lysates prepared after incubating HL-1 cells with mentioned IgG samples for 24 hours. A supernatant fraction represents non-cytoskeleton bound proteins and a pellet fraction represents cytoskeleton bound proteins. F/R (Forskolin/Rolipram) treatments served as a positive experimental control for the activation of p38MAPK. **D**. Western blot quantification of pp38MAPK in the supernatant fractions of the blots shown in A. Ratio between pp38MAPK and total p38MAPK was normalized to IgG-Control (IgG) and means represented ± SEM. N=4-6. * Indicate p<0.05, ** indicates p<0.005 compared to healthy control IgGs (IgG); # indicates p<0.05 compared to control (Ctrl) and NS is not significant. One-way ANOVA with Holm-Šídák’s multiple comparisons test was performed between IgG samples, and a student’s *t*-test was performed between control and F/R. **E.** Dissociation assays with HL-1 cells were treated in the presence or absence of 100 µM SB2021890 (SB20), which was added 30 minutes before 24-hour treatment with AC patient IgGs. Data represents the mean ±SEM. N=4. * Indicates p<0.05, SB20 compared to vehicle (DMSO) treatment. Multiple unpaired *t*-tests with Welch correction were performed.

**Fig 6:**
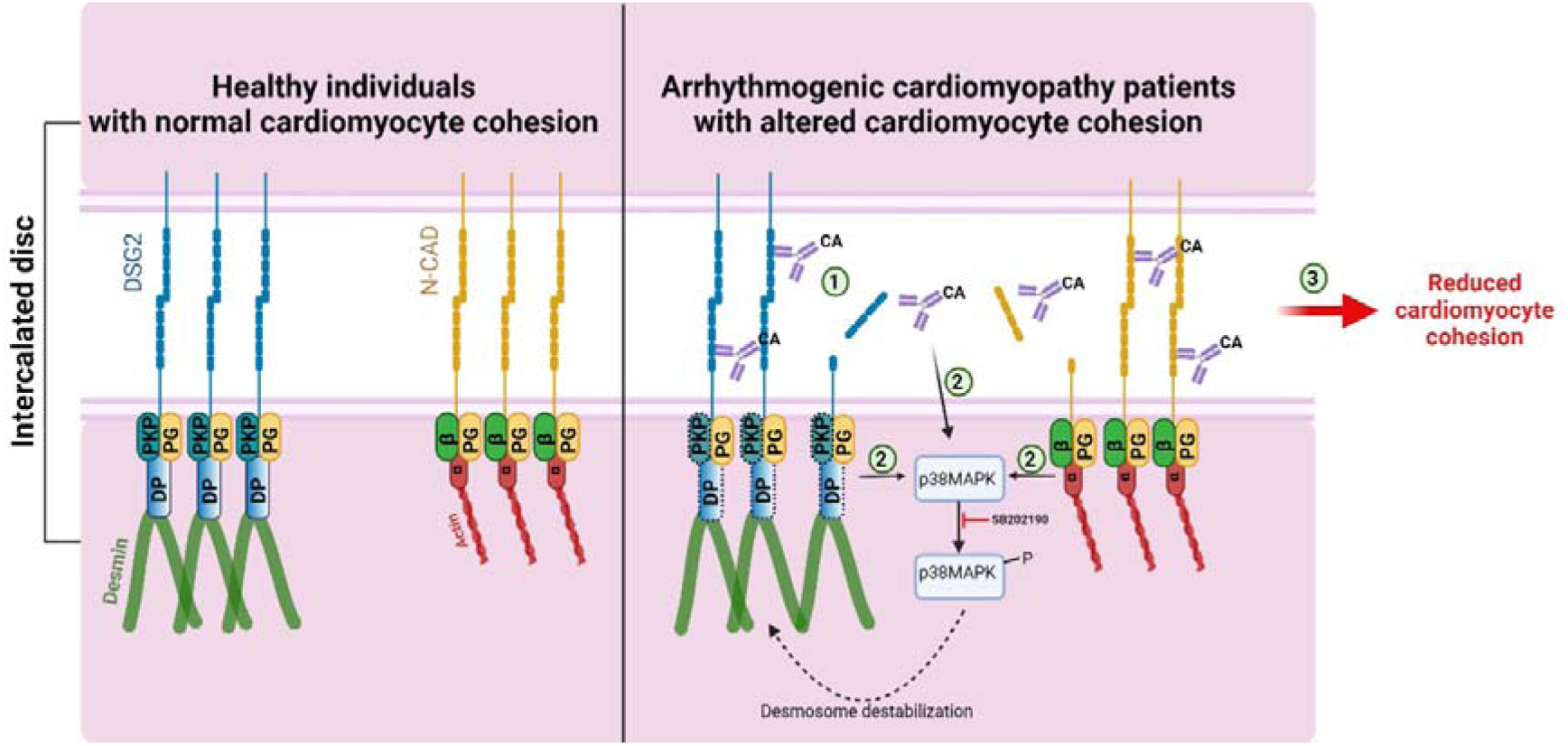
Catalytic antibodies mediated cardiomyocyte cohesion in AC patients. In normal healthy individuals the cadherin family proteins, desmoglein 2 (DSG2) and N-cadherin (N-CAD) proteins are properly localized to the ICD via the plaque proteins DSP and PKP2 and armadillo proteins PG, α-catenin (α), β-catenin (β) and YAP (Y) respectively. In AC patients, having mutations in either *DSP* or *PKP2* genes (the corresponding proteins are represented with dashed boxes), presence of catalytic antibodies (CA) that can cleave DSG2 or N-CAD could destabilize the cadherin-mediated adhesion (1), which could lead to the activation of p38MAPK (2), which further can lead to desmosome destabilization. Cleavage of cadherin proteins combined with p38MAPK activation could eventually result in reduced cardiomyocyte cohesion (3) and there by a reduction in the mechanical strengths of the cardiomyocyte. Cartoon created with BioRender.com

## Discussion

In this study, we identified for the first time that AC patient IgGs contain catalytic properties against desmosomal cadherin DSG2 and some of the tested IgGs cleave the two ICD cadherins DSG2 and N-CAD, causing loss of cadherin interaction and cardiomyocyte cohesion. Moreover, this loss of cardiomyocyte cohesion was associated with p38MAPK activation and an inhibition of p38MAPK rescued cardiomyocyte cohesion.

### Autoantibodies are prevalent in AC and occur independent of pathogenic mutations

Recently, it was reported that autoantibodies targeting DSG2 were found only in AC patients ^14^ and that anti-ICD antibodies were found in a subset of AC patients and their relatives ^15^. Since both pathogenicity and specificity of the DSG2 autoantibodies remain unclear, we characterized the effect of AC autoantibody fractions on cardiomyocyte cohesion and investigated pathogenic effects in response to the respective autoantibody binding. Immunostaining performed on human ventricular tissue detected anti-ICD antibodies within all of our 15 tested AC patients from 8 different families which is in contrast to the study by Caforio et al., in which anti-ICD antibodies were only detected in a subset of the population. This discrepancy may be due to the difference in the tissues used for detection, as the previous study used human atrium ^15^, whereas we used human ventricular tissue. Indeed, differential gene expression was observed between the human atria and ventricles in normal ^38^ and failing hearts ^39^.

### Pathogenic autoantibodies are catalytic and impair DSG2 and N-CAD homophilic interactions

We further investigated whether anti-DSG2 autoantibodies were present within AC-IgG fractions. Using ELISA, we did not detect any autoantibodies against DSG2. Chatterjee et al. used a DSG2 recombinant protein harboring the intracellular region of DSG2 and Fc-tag ^14^. In our study, we found positive signals for antibody binding to both DSG2-Fc and N-CAD-Fc following incubation with AC-IgG as well as control IgG, which did not cause ICD staining. Therefore, we assume that the false-positive signals seen in ELISA or Western blot analysis using Fc chimeric proteins were caused by a cross-reactivity of IgGs with the Fc fragment indicating that these assays cannot specifically be used to detect ICD antibodies in AC patients.

Since it is well documented that in inflammatory or autoimmune disorders, such as sepsis, HIV, autoimmune thyroiditis and myocarditis, catalytic and pathogenic autoantibodies were detectable ^28–30, 32, 33, 40–43^, we tested for the presence of catalytic antibodies in AC patients using an *in vitro* cleavage assays as described previously ^30^. We observed that 11 out 15 tested AC-IgGs cleaved the DSG2 protein containing the extracellular domains. This cleavage was completely absent in the presence of protease inhibitors, further confirming that the IgGs have protease activity. Similarly, N-CAD was also cleaved by incubation with all of the tested AC-IgG fractions..

As there are pathogenic DSG3 autoantibodies in pemphigus vulgaris and it might exhibit similar aspects of disease development and pathogenesis as AC, we performed in vitro cleavage assays utilizing DSG3, which is not expressed in cardiomyocytes. The goal of this experiment was to determine whether the observed catalytic antibodies in AC patients are specific to AC or can also be found in other desmosomal diseases such as pemphigus. Cleavage assays performed with an AC patient IgG, the monoclonal mouse PV antibody AK23 and PV patient IgG did not show any cleavage of DSG3. These results clearly established that catalytic antibodies against DSG2 and N-CAD are confined to AC patients but are absent against DSG3 in pemphigus.

Previously, it was described that DSG2 protein interaction is crucial for cardiomyocyte cohesion and that *DSG*2 gene mutations found in AC patients reduce cardiomyocyte cohesion ^19, 26, 27^. Furthermore, autoantibodies against DSG1 and DSG3 found in pemphigus cause loss of keratinocyte adhesion indicating that loss of intercellular adhesion caused by autoantibodies from patients with desmosomal diseases is a hallmark of pathogenicity^17, 37, 44, 45^. To further assess the functional relevance of catalytic antibodies, we utilized cell-free AFM force spectroscopy to measure homophilic DSG2 and N-CAD interactions. Previously, AFM experiments showed that antibodies directed against DSG2 led to a reduction in DSG2 binding *in vitro* ^27, 46^. We observed a significant decrease in DSG2 binding in response to the treatment of AC-IgG fractions which induced cleavage of ICD cadherins. In addition, AC5-IgG, apparently interfere with DSG2 binding by other mechanism such as steric hindrance which is similar to some IgGs observed in pemphigus vulgaris ^47^.

We tested whether autoantibodies in AC patients cause a loss of cardiomyocyte cohesion. Dissociation assays performed with AC patient IgGs revealed that one third of the IgGs significantly interfered with cardiomyocyte cohesion. Among the tested AC patient IgGs, 6 AC-IgG from 3 different families caused a significant decrease in cardiomyocyte cohesion in cultured cardiomyocytes. A modified dissociation assay in murine slice cultures ^26^ with AC1-IgG and AC15-IgG (from two different families and containing different gene mutations) confirmed that autoantibodies impair cardiomyocyte cohesion in intact cardiac tissue. However, since not all patients’ IgG which induced DSG2 cleavage, reduced cardiomyocyte cohesion, we speculate that the amount or activity of catalytic antibodies may vary in patients. This may not become obvious in cleavage assays with a defined concentration of recombinant protein which presumably is more accessible to cleavage compared to DSG2 and N-CAD assembled in intercellular contacts of living cells.

### Pathogenic autoantibodies induce signaling associated with loss of desmosomal adhesion

As it has been shown that pathogenic antibodies targeting DSG1 and DSG3 lead to p38MAPK activation in pemphigus ^35–37^, we investigated whether this signaling pathway is activated by AC autoantibodies. Similar to PV, we showed that in cardiomyocytes an inhibition of p38MAPK enhances cardiomyocyte cohesion ^19^. Here, we found that most AC-IgG fractions which reduced cardiomyocyte cohesion activated p38MAPK in HL-1 cells, whereas inhibition of p38MAPK rescued from the loss of cardiomyocyte cohesion. Therefore, similar to pemphigus ^35–37^, autoantibodies in AC patients are capable of activating p38MAPK consequently contributing to a loss of cardiomyocyte cohesion. In addition, the role of autoimmunity in the pathogenesis of AC is in line with recent findings that inflammation is also involved in AC pathogenesis ^48^.

In conclusion, our data demonstrates that in AC patients anti-ICD antibodies are frequent, and contain catalytic antibodies against DSG2 and N-CAD, which interfere with cadherin interaction and activates p38MAPK and thereby contribute to a loss of cardiomyocyte cohesion.

## Methods

### IgG preparation

After collecting the blood samples, IgGs from patients suffering from AC or PV were prepared as described previously ^37^. AK23, a monoclonal pathogenic antibody, derived from a pemphigus mouse model, was purchased from Biozol (Eching, Germany).

### Collection of human left ventricle tissue and immunostaining

Human left ventricle tissue of a female was collected post-mortem, embedded in Richard-Allan Scientific Neg 50 Frozen Section Medium (Thermo Scientific, #6502G;), snap frozen in liquid nitrogen, and cut into 7 µm thin sections with a cryostat (Thermo Scientific, Cryostar NX70). Sections were then heated to 37°C for 8 minutes, washed in PBS 3 times and fixed in 2 % PFA for 10 minutes, and washed 5 minutes again in PBS 3 times. Then sections were permeabilized with 1 % Triton-X-100 for 60 minutes, washed in PBS and treated with PBS with 50 mM ammonium chloride (NH_4_Cl buffer) for 20 minutes. After washing with PBS, the sections were blocked with 3 % bovine serum albumin and 10 % normal goat serum in PBS for 60 minutes. Then the sections were incubated either with respective IgGs diluted 1:50 in PBS, or polyclonal rabbit anti-DSG2 (Progen, #610121), and monoclonal mouse anti-N-CAD (BD Transduction, #610921) antibodies at 4°C overnight. Sections were washed with NH_4_Cl buffer for 5 minutes 3 times, and Alexa Fluor 488-conjugated goat anti-mouse secondary antibodies (Dianova, #115-545-062) and Cy3-conjugated goat-anti-human (Dianova, #109-165-008) along with DAPI (Roche, #10236276001) at 1:2000 were incubated for 60 minutes at room temperature. Then, sections were washed for 5 minutes 3 times again. ProLong™ Diamond Antifade Mountant (Thermo Scientific, #P36961) was used to mount the samples. Images were acquired with the Leica SP5 confocal microscope equipped with a 63×□ NA 1.4 PL APO objective using LAS-X software (Leica Microsystems, Wetzlar, Germany).

### Cell cultures, dissociation assays, and transient transfection

#### HL-1 cells

The murine atrial cardiac myocyte cell line HL-1 was maintained as explained previously ^19^. In brief, cells were grown in Claycomb medium (#51800C) supplemented with 10 % fetal bovine serum (#F2442), 100 µmol L^-1^ norepinephrine, 100 µg ml^-1^ penicillin/streptomycin, and 2 mmol L^-1^ L-glutamine at 37°C, 5 % CO_2_ and 100 % humidity as described previously. All cell culture reagents were purchased from Sigma-Aldrich, Munich, Germany. After seeding for experiments, cells were incubated in Claycomb medium without norepinephrine to avoid basal adrenergic stimulation.

### Dissociation assays in HL-1 cells

For dissociation assay experiments, cells were seeded at 125000 cells per cm^2^ on cell culture plates coated with 0.02 % gelatin and 25 µg ml^-1^ fibronectin, grown for 72 hours and then treated with respective IgGs for 24 hours. Subsequently, the cells were washed with HBSS and treated with Liberase-DH (0.065 U.ml^-1^, Sigma-Aldrich Munich, #5401054001) and Dispase II (2.5 U.ml^-1^, Sigma-Aldrich, #D4693-1G), and incubated at 37°C until the cell monolayer detached from the wells. Then, the enzyme mix was carefully removed from the wells and replaced by HBSS. Mechanical stress was applied by horizontal rotation on an orbital shaker (Stuart SSM5 orbital shaker) at 1.31 g for 10-15 minutes. The number of fragments were determined by counting utilizing a binocular stereomicroscope (Leica microsystems, Wetzlar, Germany).

### Dissociation assays in murine cardiac slices

Twelve weeks old age and sex-matched littermate of wild-type mice were used for experiments. mice were sacrificed by cervical dislocation, hearts were immediately placed in pre-cooled oxygenated cardiac slicing buffer, embedded in low melt agarose, and 200 µm thickness slices were cut with a LeicaVT1200S vibratome (Leica Biosystems). For dissociation assays, slices were washed gently with HBSS, transferred to pre-warmed cardiac slices medium, and incubated for 1 hour with indicated mediators at 37°C, 5 % CO_2._ Dissociation assays in murine cardiac slices were performed similarly to dissociation assays in HL-1 cells. However, Liberase-DH and Dispase II were added together for 30 minutes in this case. Next, MTT was added and mechanical stress was applied using an electrical pipette by pipetting up and down. Finally, the same mechanical stress was applied to the other slices of the same experiment. Then the content of each well was filtered using a 70 µm nylon membrane, and pictures of the well were taken and stitched together using the AutoStich software. Only the number of dissociated cardiomyocytes stained with MTT were counted for quantification. To control variations due to different slice size or location in the ventricle, consecutive slices for control and treatment were used, respectively, and the result of a slice was normalized to the respective control slice.

### ELISA

#### Using DSG2-His protein

The cDNA of the extracellular domain of human DSG2 [NP_001934.2], was added with His-tag and inserted into pcDNA3.4 TOPO vector by pcDNA3.4 TOPO TA cloning kit (Thermo Fisher Scientific, MA) according to the manufacturer’s protocol. Subcloned plasmids were used to transform into Expi293F cells (Thermo Fisher Scientific, # A14527) using the Expi293 expression system (Thermo Fisher Scientific, # A14635) in an expression medium for 5 days as per manufacturer’s instructions with D-PBS (+) preparation reagent (Ca^2+^, Mg^2+^ solution; Nacalai tesque, Kyoto, Japan) added one day after transformation. The proteins were purified from the supernatant using affinity Ni^2+^ charged resin chips (PS Tips IMAC, Mettler Toledo, OH) and buffer were then replaced into Tris-buffered saline (TBS) with 1 mM CaCl_2_ (TBS-Ca). The solution was adjusted to 5 µg ml^-1^ in a total of 100 µl per well diluted antigen and were coated into Pierce™ Nickel Coated Plates (Thermo Fisher Scientific, #15142) at 4°C overnight. After the coating solution was removed, the wells were washed twice with PBS (pH 7.4). Samples (AC patient IgGs and control IgGs) were diluted 1:10 and 1:100 in blocking buffer and each of the sample was incubated for 1 hour at room temperature. As positive control, anti-human DSG2 antibodies (Abcam, #ab14415) were used at 10 µg ml^-1^ dilutions in blocking buffer. After washing 4 times with PBS, the wells were incubated with either goat anti-mouse IgG-HRP (Cell signaling #91196) at 1:2000 dilution in blocking buffer for DSG2-Abcam antibody, or anti-human IgG-HRP (from the kit, MBL MESACUP™-2 TEST Desmoglein3, #7885E) for the patient samples for 1 hour. Finally, the wells were washed 4 times with blocking buffer and incubated with 100 µl TMB substrate solution (from the Kit, MBL MESACUP™-2 TEST Desmoglein3, #7885E). After 30 minutes incubation at room temperature, 50 µl TMB stop solution (from the kit, MBL MESACUP™-2 TEST Desmoglein3, #7885E) was added and the OD was measured at 450 nm using a microplate reader (Bio-Rad Laboratories, Hercules, CA, U.S.A.).

#### Using DSG2-Fc protein

ELISA using DSG2-Fc protein was performed according to the protocol from Chatterjee et al. with adaptations ^14^. First, Nunc MaxiSorp™ flat-bottom plates (Thermo Scientific; # 44-2404-21) were coated with 2 µg ml^-1^ recombinant human DSG2 (Creative Biomart; #DSG2-1601H) in 100 mM bicarbonate/carbonate coating buffer (3.03 g Na2CO3; 6.0 g NaHCO3; ad. 1000 ml distilled water; pH 9.6). 100 µl per well diluted antigen was used and incubated for 2 hours at room temperature. For each incubation step, the plate was covered with adhesive plastic films. Subsequently, after the coating solution was removed, the wells were washed twice with PBS (pH 7.4) and then incubated with 100 µl blocking buffer (PBS with 2 % BSA) for 2 hours at room temperature. Samples (patient IgGs and control IgGs) were diluted 1:50 in blocking buffer, and 100 µl each of the samples was incubated overnight at 4°C. As a positive control, two different anti-human DSG2 antibodies (DSG2-Origene, #BM5016; DSG2-Abcam, #ab14415) were used at 1:1000 dilutions in blocking buffer. The next day, after being washed 4 times with PBS, the wells were incubated with either goat anti-mouse IgG-HRP (Dianova, #115-035-068) at 1:2000 dilution in blocking buffer for DSG2-Origene antibody and DSG2-Abcam antibody, or goat anti-human IgG-HRP (AffiniPure F(ab’)□ Fragment Goat Anti-Human IgG (H+L) Pox; Jackson Immuno Research # 109-036-088) at 1:100000 dilution in blocking buffer for the patient samples. Finally, the wells were washed 4 times with blocking buffer and incubated with 100 µl TMB substrate solution (Thermo Scientific; #N301). After 20 minutes of incubation at room temperature, 100 µl TMB stop solution (Thermo Scientific; #N600) was added, and the OD was measured at 450 nm using a Tecan plate reader.

### Western Blot to detect autoantibodies against DSG2

Western blot was performed to detect anti-DSG2 autoantibodies, based on the methods reported by Chatterjee et al. ^14^ with adaptations. Recombinant protein of human DSG2 tagged with IgG Fc domain (Fc) (Creative Biomart; #DSG2-1601H) and N-CAD-Fc (Sino Biological; #11039-H03H) was reconstituted according to manufacturer’s instructions and diluted in distilled water to a concentration of 100 ng µl^-1^ or 250 ng µl^-1^, respectively. For SDS-polyacrylamide gel electrophoresis, 100 ng of either DSG2 or N-CAD were mixed with Laemmli sample buffer containing DTT, boiled for 5 minutes at 95°C and loaded onto a 7.5 % polyacrylamide gel with stacking gel. The gel electrophoresis was run in a Bio-Rad Mini-PROTEAN® Tetra Cell system, by starting with constant 80 V for 20 minutes, followed by constant 120 V for additional 45 minutes. For molecular weight determination, 5 µl of Page Ruler Plus™ (Thermo Scientific; #26619) was also run on the same gel. The separated proteins were transferred onto a 0.45 µm nitrocellulose membrane (Thermo Scientific; #LC2006) for 90 minutes at a constant 350 mA using Bio-Rad Mini-PROTEAN® Tetra System. To block non-specific binding sites, the membranes were incubated at room temperature for 1 hour in TBS with 0.1 % Tween 20 (TBST) containing 5 % non-fat dry milk (Sigma Aldrich; #70166-500G). The membranes were incubated with AC patient IgGs, as well as pooled isolated IgG taken from four healthy donors (2 males and 2 female) as a control, diluted 1:50 in 5 % BSA/TBST overnight at 4°C (BSA: VWR; #422361V). As a negative control, only secondary antibody was used. Membranes were then washed extensively 3 times in TBST for 15 minutes and incubated in goat anti-human-HRP conjugated secondary antibody (Jackson Immuno Research; #109-035-003) at 1:20000 dilution in TBST for 1 hour at room temperature. After incubation, the membranes were washed 3 times for 10 minutes in TBST and immunoreactive bands were detected with ECL solution SuperSignal™ West (Thermo Scientific; #34577) after 40 seconds of exposure in a FluorChemE system (ProteinSimple California, USA)

### DSG2 and N-CAD cleavage assay

IgGs from AC patients (AC1-AC15), or a PV patient IgG and murine pemphigus monoclonal antibody AK23 and IgGs pooled from 4 healthy control donors (2 male, 2 female; referred as IgG) were incubated for 10 minutes at room temperature with or without protease inhibitors (Roche, cOmplete™: #11697498001; cOmplete™, EDTA-free: #11873580001) in buffer A (10 mM HEPES, pH 7.5; 0.02 % NaN3; 0.05 % Brij 35) containing DMSO (1 % v/v) in the final reaction mixture. After the incubation, 200 ng of hDSG2-Fc ^49^, 200 ng of hN-CAD-Fc (#11039-H03H) or hDSG3-Fc ^47^ [all DSG-Fc constructs include the complete extracellular domain and Fc-Fragment of human IgG1] were added to a final reaction volume of 10 µl and incubated for 4 hours at 37°C. Finally, the results were analyzed by SDS-PAGE. DSG2 and DSG3-Fc proteins and N-CAD-Fc protein were detected with the following antibodies: mouse-anti-DSG2 (#BM5016, Origene, 1:200); mouse-anti-DSG3 5G11 (#32-6300, Invitrogen, 1:500) and N-CAD-Fc protein with (#66219-1-Ig, proteintech; 1:200).

### Atomic force microscopy (AFM)

Atomic Force Microscopy measurements were performed as described before ^26, 50^ using a Nanowizard® III AFM (JPK Instruments, Berlin, Germany) with an optical fluorescence microscope (Axio Observer D1, Carl Zeiss) at 37°C. AFM cantilevers (MLCT AFM Probes, Bruker, Calle Tecate, CA, USA) were coated with recombinant DSG2-Fc or N-CAD-Fc as described previously ^51^. Force-distance curves (FDC) were recorded with a loading force of 0.3 nN, a contact delay of 0.1 s and a retraction velocity of 1 µm s^-1^. For each individual experiment, 1000 FDC were recorded in HBSS buffer to obtain the baseline trans-interaction probability. Thereafter, the mica and cantilever were incubated for 1 hour with respective AC patient IgGs (diluted 1:50) or control-IgG (diluted 1:50) and 1000 additional FDC were recorded with the same settings and on the same position on the mica sheet. JPKSPM Data Processing software (JPK Instruments) was used to evaluate acquired FDC. Characteristic unbinding signatures in the FDC were analyzed to calculate an interaction probability for each condition. To account for variations in cantilever and mica sheet coating the binding probability was normalized to the baseline binding probability for each experiment.

### Protein extraction for Western blotting with triton buffer

To separate the soluble cytosolic fraction from the insoluble cytoskeletal or membrane-bound fraction protein, extraction with triton buffer (0.5 % Triton-X-100; 50 mM MES; 25 mM EGTA; 5 mM MgCl2; pH 6.8) was performed. The triton extraction buffer was supplemented with a cOmplete™ Protease Inhibitor Cocktail (Roche; #11697498001) and PhosSTOP™ (Roche; #4906845001). The cells in a 12-well plate were washed twice with ice cold PBS and incubated with 70 µl supplemented triton extraction buffer for 20 minutes on ice with gentle movement. After incubation, all cells were scraped and transferred into a tube, and soluble and pellet fractions were separated by centrifugation for 5 min at 4°C with 15000 g in an Eppendorf 5430R centrifuge. The supernatant was then transferred into a clean tube and marked as soluble fraction. The pellet was washed with triton extraction buffer and lysed in 60 µl lysis buffer (12.5 mM HEPES; 1 mM EDTA disodium; 12.5 mM NaF and 0.5 % SDS; pH 7.6) supplemented with cOmplete™ Protease Inhibitor Cocktail and PhosSTOP™ and was finally sonicated for homogenization.

### Statistical analysis

Statistical analysis was performed using GraphPad Prism 9 (GraphPad Software, La Jolla, USA) as described in the corresponding figure legend. Significance was assumed at P < 0.05.

### Study approval

Blood samples were collected from AC patients and healthy donors under the Ethics Committee’s approval at Ludwig-Maximilians-University (LMU), Munich, Germany (Project no: 20-1061), and informed written consent was obtained. In addition, human heart ventricular tissues were collected from body donors at the Institute of Anatomy and Cell Biology, LMU, Munich, Germany. Written informed consent was obtained from the female body donor for the use of tissue samples in research. The body arrived within 17 hours after decease and was free of any cardiac disorders. The pemphigus patient gave written consent for research use. A positive vote of the Ethics Committee from the Medical Faculty of the University of Marburg (Az 20/14) was given.

Animal handling, breeding, and sacrifice were performed under approval of regulations by the regional government Upper Bavaria (Gz. ROB-55.2-2532.Vet_02-19-172).

## Author contributions

S.Y., K.S., A.J., M.H^1^., M.S., D.K., S.E., I.N, and K.H., acquired and analyzed the data, T.H., and D.S. analyzed the data, S.Y. and J.W. drafted the manuscript. S.Y., K.S., T.H, D.S., S.K., and J.W. made critical revision of the manuscript for important intellectual content, M.H^2^., M.W., B.M.B., R.B., and S.K. provided reagents, S.Y. and J.W. handled supervision and designed the research.

## Acknowledgments

We thank Martina Hitzenbichler, and Kilian Skowranek for technical assistance.

## Data availability statement

The data underlying this article will be shared on reasonable request to the corresponding author.

