## Supplementary file for "Catalytic antibodies in arrhythmogenic cardiomyopathy patients cleave desmoglein 2 and N-cadherin and impair cardiomyocyte cohesion"

**Supplementary figures.**

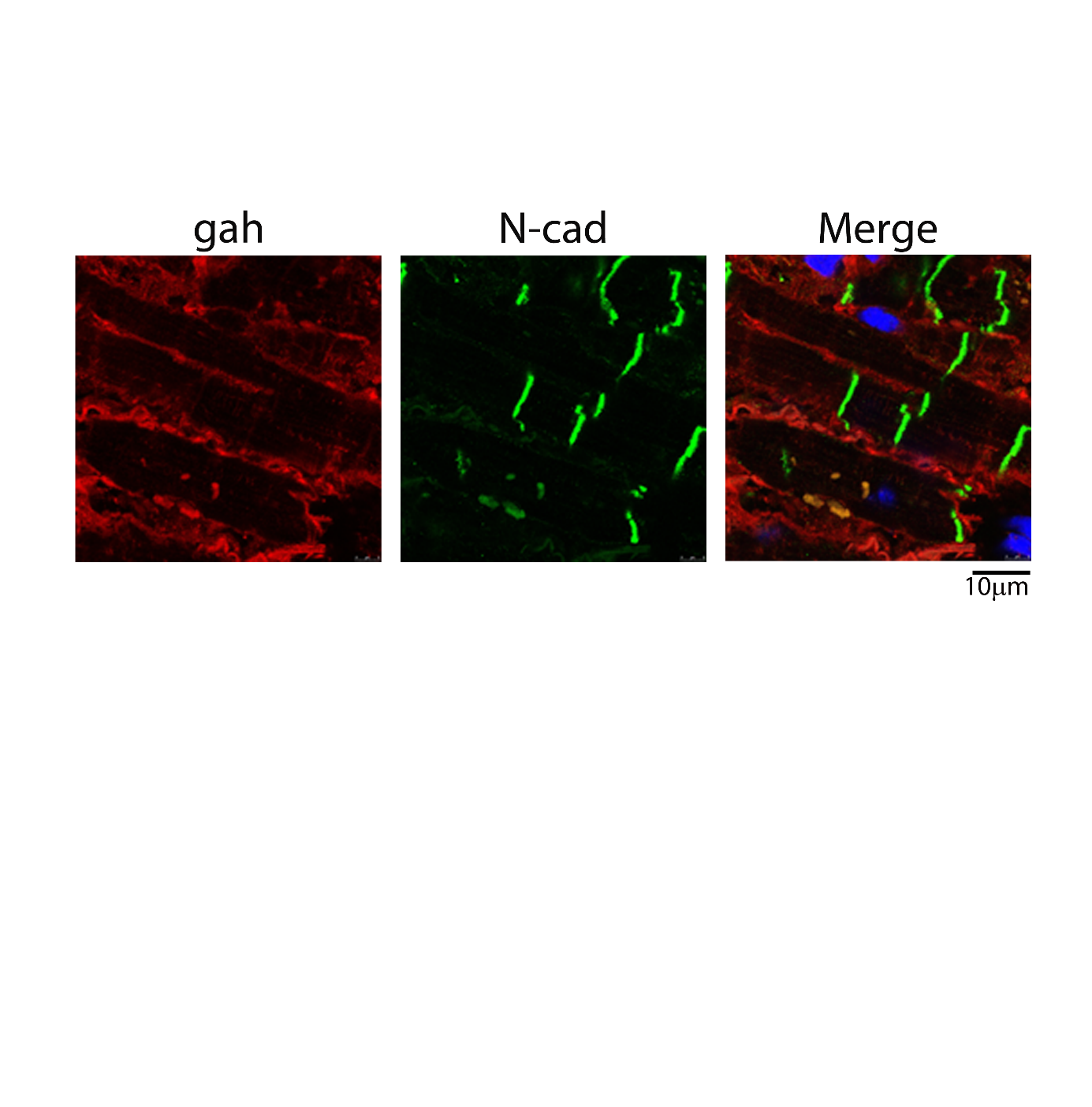

**Supplementary figure 1:**

Human left ventricular tissue was incubated with goat anti-human tagged with Cy3. N-cadherin (N-CAD) was used as a marker for ICD. The image shows no positive staining of ICD. Images represent immunofluorescence analysis of 3 repeats. Scale bar:10 µm.

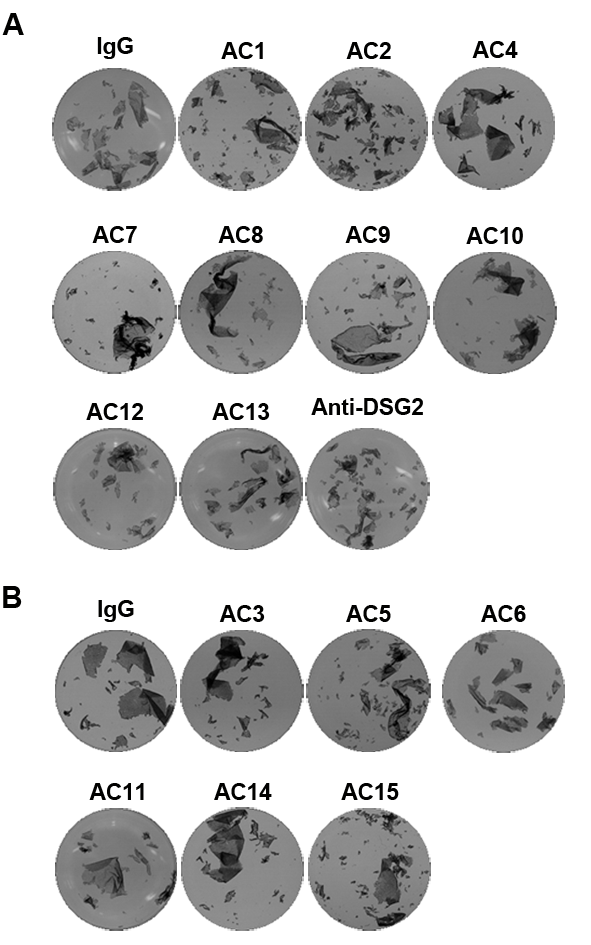

**Supplementary figure 2:**

Dissociation assays were performed in HL-1 cells treated with respective IgGs, as was used in figures 2 B and C. Representative images of fragments obtained after treatment and dissociation assays.

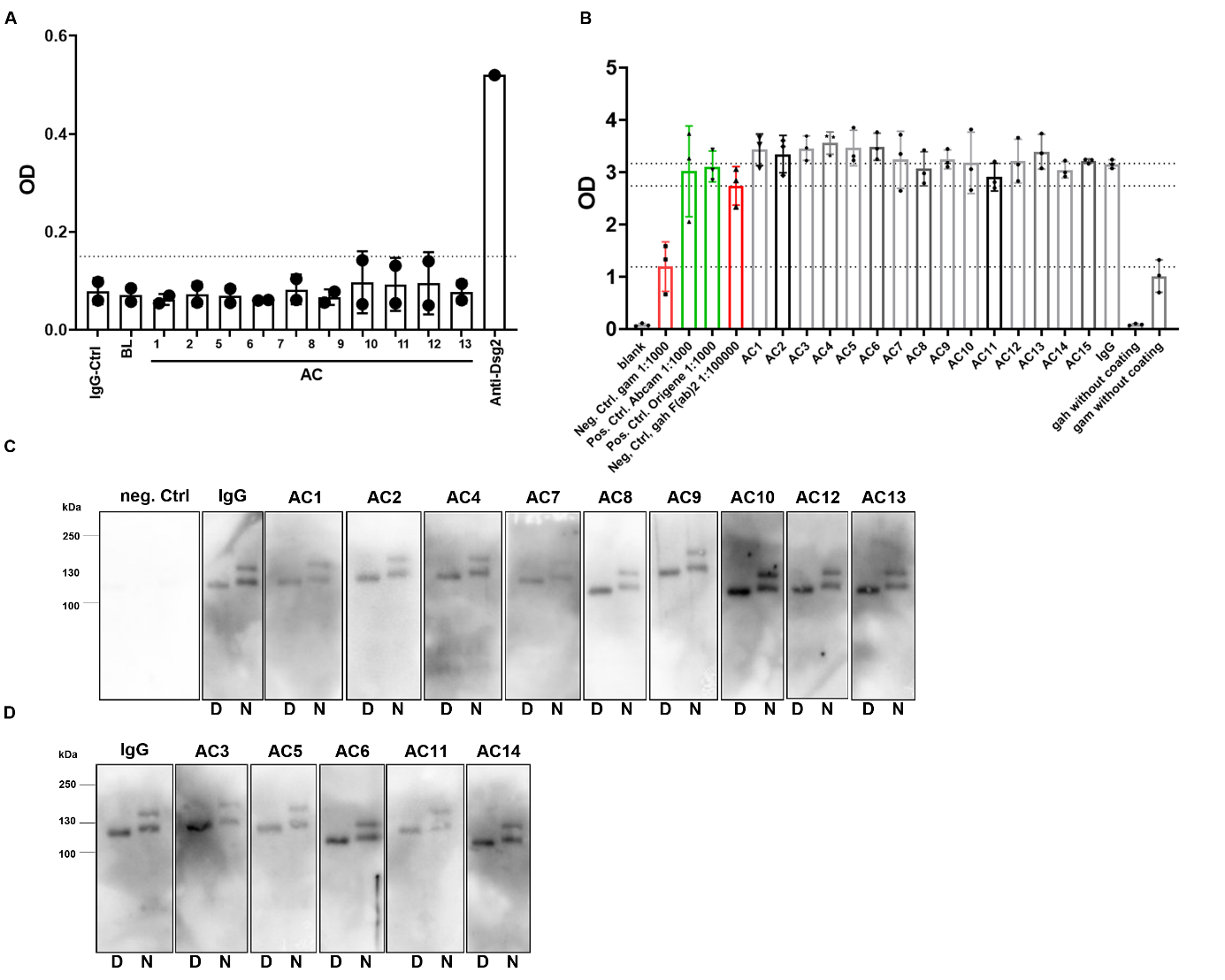

**Supplementary figure 3:**

**A.** ELISA analysis of AC patient IgG samples (diluted 1to10) for autoantibodies against human DSG2 protein. Y-axis represents optical density (OD) measured at 450 nm. An OD value of 0.15 was considered a cut-off value, as represented in the bar graph with the dotted line. Bar graphs represent mean values ± SD. N=2. **B.** ELISA analysis of AC patient IgGs (diluted 1to50) for autoantibodies against human DSG2-Fc protein. Y-axis represents optical density measured at 450 nm. In the bar graph, dotted lines were drawn to compare with respective positive and negative controls. Bar graphs represent mean values ± SEM. N=3. **C,D.** Blotting of 100 ng human DSG2 and N-CAD fusion protein (with Fc), incubated with healthy control-IgG (IgG), DPC patient IgG (**C**) and ARVC patient IgG samples (**D**), with goat-anti-human IgG-HRP as a secondary antibody. Representative image of Western blot analysis performed in 3 experimental repeats.

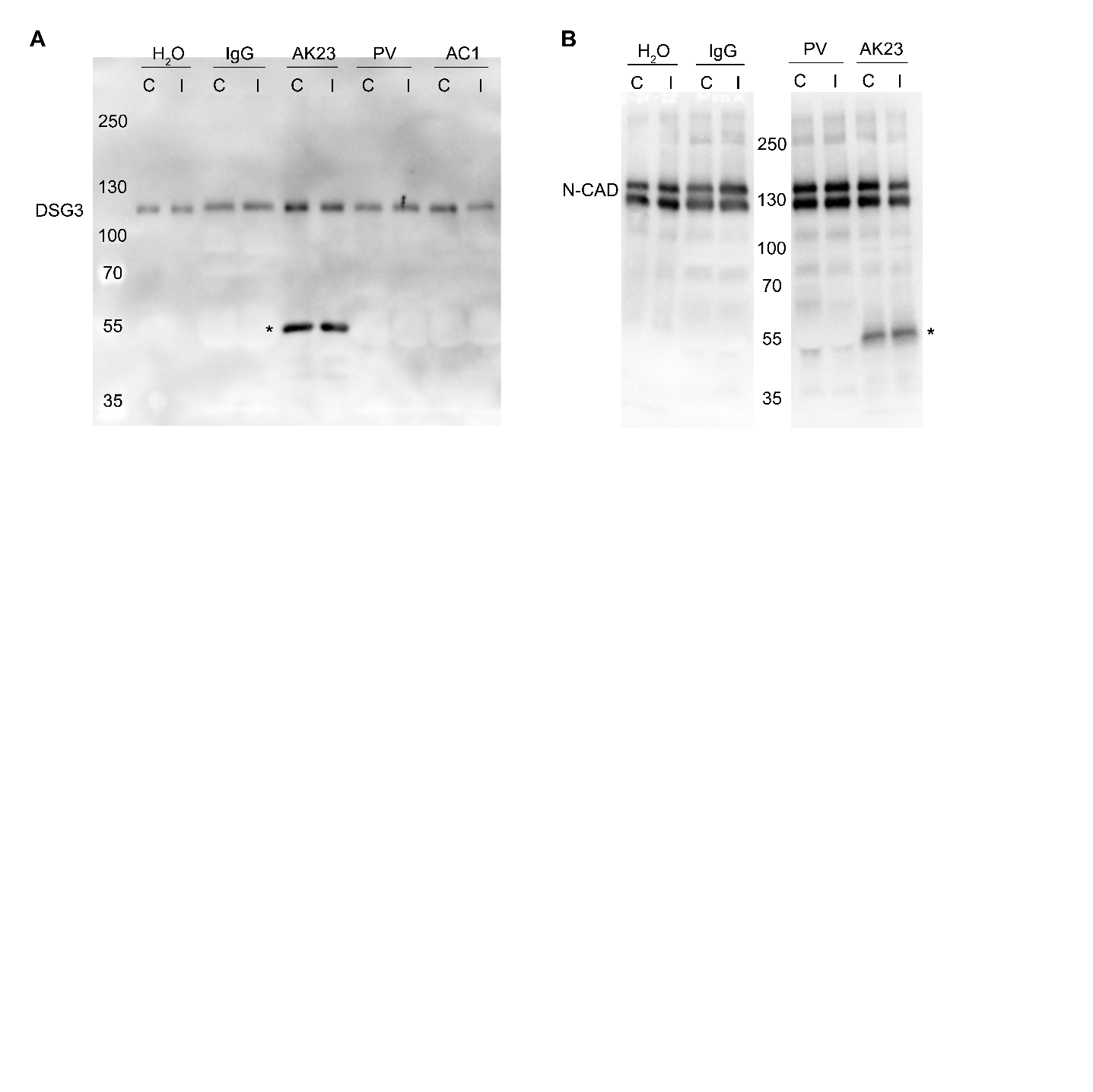

**Supplementary figure 4: DSG3-Fc cleavage assay and N-CAD-Fc cleavage with PV and AK23**

**A,B**. DSG3-Fc protein (**A**) or B. N-CAD-Fc protein (**B**) was incubated with the respective IgGs as shown in the figure, with and without protease inhibitors for 4 hours. AK23 and pemphigus vulgaris (PV) IgGs were used as controls that contain anti-DSG3 antibodies. Representative image of Western blot analysis performed in 3 experimental repeats.

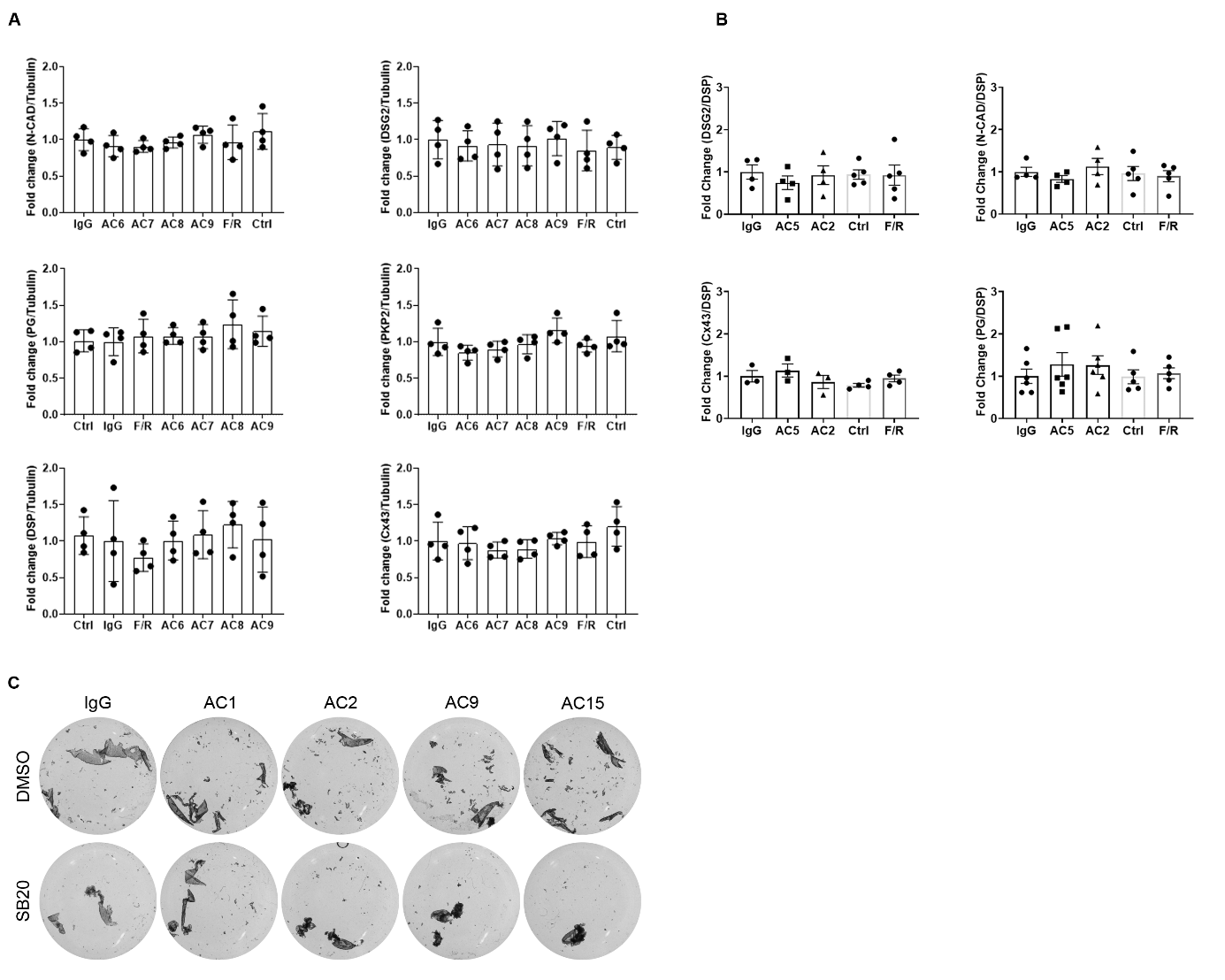

**Supplementary figure 5:** Western blot quantification of proteins shown in figure 5.

**A**. Western blot quantification of the respective proteins for the blots shown in Figure 5A. α-Tubulin (Tubulin) was used as internal control and mean values were normalized to healthy control IgG. One-way ANOVA with Dunnett post-hoc was performed between IgG samples and a Student’s-t-test was performed between control and F/R. N= 4. **B**. Western blot quantification of proteins in the pellet fractions of the blots shown in figure 5C. DSP was used as an internal control for pellet fractions (non-cytoskeleton bound fractions) and mean values were normalized to IgG-Control. (IgG). * indicate p<0.05, One-way ANOVA with Dunnett post-hoc was performed. N=3-5. **C**. Representative images of fragments obtained after treatment and dissociation assays performed for figure 5E.

Supplementary Table 1: Details of antibodies used in this manuscript.

| **Name of the antibody** | **Company and catalog number** | **Dilution used for WB** | **Dilution used for IF** |
| --- | --- | --- | --- |
| phospho p38 | Cell Signaling, #4511 | 1:1000 |  |
| p38 | Cell Signaling, #9211 | 1:1000 |  |
| N-Cadherin | BD Transduction, #610921 | 1:1000 | 1:500 |
| Demoglein 2 | Progen, #610121 | 1:500 | 1:300 |
| Desmoplakin | Progen, #61003 | 1:500 |  |
| Plakoglobin | Progen, #61005 | 1:1000 |  |
| Plakophilin2 | Progen, #651167 | 1:25 |  |
| Cx43 | Sigma#SAB4501175 | 1:1000 |  |
| Goat anti-mouse Alexa-488 | Dianova #115-545-003 |  | 1:600 |
| Goat anti-rabbit Cy5 | Dianova,#111-175-144 |  | 1:600 |
| Goat-anti-human HRP (H+L) | Jackson Immuno Research  #109-035-003 | 1:20000 |  |
| Goat-anti-human F(ab)_2_ HRP (H+L) | Jackson Immuno Research  #109-036-088 | 1:100000 for ELISA |  |
| Goat-anti-human Cy3 | Dianova#109-165-008 |  | 1:600 |
| Tubulin | Abcam# Ab7291 | 1:4000 |  |

WB; Western blot. IF; immunofluorescence study

Supplementary Table 2: Details of mediators used in this manuscript.

| **Name** | **Company** | **Catalog.No** | **Concentration** |
| --- | --- | --- | --- |
| Forskolin (F) | Sigma-Aldrich | #F3917 | 5 µmol/L |
| Rolipram (R) | Sigma-Aldrich | #R6520 | 10 µmol/L |
| SB202190 (SB20) | Cell signaling | #8158S | 100 µmol/L |
